# MEG Signatures of Differences in Auditory Cortex Responses to Trigger versus Non-Trigger Sounds in Misophonia

**DOI:** 10.64898/2026.09.14.751161

**Authors:** Jasmine Tan, Sergio Osorio, Grace Levine, Zein Nayal, Seppo P Ahlfors, Julie G. Arenberg, Nathaniel Mercaldo, Jyrki Ahveninen, Tommi Raij, Tal Kenet

## Abstract

Misophonia is characterized by intense emotional and physiological reactions to specific “trigger” sounds that may vary across individuals. Despite greater recognition and increased research, the neural mechanisms underlying misophonia are still poorly understood. Using magnetoencephalography (MEG) with millisecond temporal resolution, we examined auditory cortex responses to trigger versus non-trigger sounds in 23 individuals (ages 14 – 32 years) that met criteria for misophonia on a screening assessment. We found that in the right hemisphere, trigger sounds elicited significantly smaller evoked responses, lower power of induced alpha-band oscillations, and stronger alpha-gamma cross-frequency phase-amplitude coupling, compared to non-trigger sounds. We also found that evoked responses correlated with behaviorally assessed extent of misophonia symptoms. These findings support the idea that misophonia is characterized at least in part by altered neurophysiological processing of auditory trigger stimuli early in the auditory cortex hierarchy. Our results provide novel insight into the neural basis of misophonia and underscore the importance of examining brain activity even at the earliest levels of the cortical hierarchy with high spatio-temporal resolution.

## 1. Introduction

Misophonia is a condition in which specific sounds elicit intense, disproportionately negative emotional and physiological reactions, including disgust, rage, and anxiety (Schroder et al 2013, Wu et al 2014). The trigger sounds in misophonia are typically repetitive and of human or biological origin — such as chewing, lipsmacking, breathing, or keyboard clicking — and are perceived as innocuous by individuals without the condition (Hansen et al 2021). Misophonia is increasingly recognized as a distinct clinical entity that is associated with significant functional impairment, including social withdrawal, avoidance behaviors, and reduced quality of life (Rosenthal et al 2023, Rosenthal et al 2022, Rosenthal et al 2026, Swedo et al 2022, Wu et al 2014).

Despite growing clinical recognition, misophonia remains incompletely characterized at both behavioral and neurobiological levels. Early theoretical accounts framed misophonia as a conditioned aversive reflex, whereby neutral sounds acquire threat salience through associative learning processes, implicating limbic structures such as the amygdala (Jastreboff & Jastreboff 2023). Neuroimaging studies have since confirmed heightened amygdala activation and increased functional connectivity between auditory cortex and limbic structures in misophonia (Kumar et al 2017), and resting-state fMRI work has revealed atypical connectivity involving the anterior insular cortex, suggesting a role for interoceptive and salience processing networks (Eijsker et al 2019, Schroder et al 2019). Nonetheless, the neural correlates of misophonia remain poorly understood, particularly at the level of early sensory processing areas. For instance, it is unclear whether misophonic reactions arise primarily from downstream limbic and evaluative processes or whether they are preceded by, or even dependent upon, alterations in the earliest stages of auditory cortical processing.

To date, most prior neuroimaging work has employed functional MRI (Kumar et al 2021, Kumar et al 2017), which offers excellent spatial but poor temporal resolution. Thus, relatively little is known about the spectrotemporal dynamics of early auditory cortex responses to trigger versus non-trigger sounds. This is a significant gap, because the auditory cortex actively shapes perception on a timescale of milliseconds (Giraud & Poeppel 2012). For instance, alpha-band (8–12 Hz) oscillations in the auditory cortex are thought to reflect inhibitory gating mechanisms that regulate the flow of sensory information (Klimesch 2012), while gamma-band (>30 Hz) activity has been linked to active cortical processing and feature binding (Fries 2015). Furthermore, the coupling between slower and faster oscillatory rhythms, commonly known as cross-frequency coupling (CFC), has emerged as a key candidate mechanism through which hierarchical cortical computations may be organized (Canolty & Knight 2010). Mapping these dynamics requires methods with high temporal resolution, such as magnetoencephalography (MEG) or electroencephalography (EEG). Understanding whether and how trigger sounds modulate early auditory cortical responses is therefore an essential part of constructing a complete account of the neural basis of misophonia, and may reveal mechanisms that are invisible to slower hemodynamic imaging methods.

For this study, we tested whether responses in the auditory cortex differ for trigger versus non-trigger sounds in individuals with misophonia. We hypothesized that if misophonia involves altered early auditory processing, then auditory cortex dynamics should differ between sound categories from the early stages of cortical response. To test our hypotheses, 23 participants with clinically confirmed misophonia underwent MEG recording while listening to individually tailored trigger sounds and matched non-trigger control sounds of comparable acoustic complexity and overall amplitude. Source reconstruction was used to extract evoked responses from the auditory cortex bilaterally, and time-frequency analyses were applied to characterize oscillatory responses in the alpha, beta, and gamma bands. We additionally computed phase-amplitude CFC to probe the organization of oscillatory hierarchies within the auditory cortex. By focusing on auditory cortex activity within the first few hundred milliseconds post-stimulus onset, we aimed to delineate early processing stages that may act as the initial catalyst for downstream emotional and autonomic symptoms characteristic of the condition.

## 2. Methods

### 2.1 Participants

While misophonia is not currently listed in the DSM-5, it is becoming increasingly recognized as a distinct condition, and several behavioral assessments have been developed and validated to screen for it. Because it is not yet clinically recognized, we began by recruiting participants with self-reported misophonia. Out of 72 screened participants, 40 (35 females, 3 males) met the criteria for misophonia as measured by the Selective Sound Sensitivity Syndrome Scale, known as the S-Five (Vitoratou et al 2021), and were recruited for the study. After 17 of the datasets were collected, an error was found in the normalization of the trigger versus non-trigger sounds, resulting in different overall loudness levels. Out of an abundance of caution, data here are presented from the 23 participants (20 females, 3 males) that had their data collected after the error was fixed, and the sounds correctly normalized. All participants had IQ ≥ 85 as measured by the Kaufman Brief Intelligence Test – II (KBIT-2, Kaufman & Kaufman 2004), were right handed as confirmed using an updated Edinburgh handedness survey (Oldfield 1971), and ranged in age from 14 to 32 (mean (s.d.): 23.8 (5.0)). Informed consent was obtained from all participants according to protocols approved by the Institutional Review Board of Mass General Brigham (MGB).

### 2.2 Audiological Assessments

All participants underwent a standard audiology assessment battery at Massachusetts Eye and Ear of MGB. The assessment battery consisted of: (1) Comprehensive audiogram, with extended high frequencies; (2) Auditory Brainstem Responses; and (3) Otoacoustic Emissions (OAEs). All participants had normal hearing, based on pure tone thresholds, at thresholds less than or equal to 25 dB HL through air and bone conduction transducers at all frequencies (250, 500, 1000, 2000, 4000, 8000 Hz). One participant had an abnormal ABR with the right ear having prolonged latencies of wave I and V, with no discernible wave III. All other audiometric data were normal for this participant.

### 2.3 Paradigm

Participants were performing an auditory spatial attention task while MEG data were collected. The MEG recording session consisted of 6 runs of about 5 min each, with as many breaks as needed between the runs. Each run consisted of 8 blocks, and each block of 36 trials began with participants visually cued with an arrow (duration 2 seconds) to attend to the right or left ear. In total, per participant, we collected an average of 78 trigger trials (SD = 6.5, 64-85) and 75 non-trigger trials (SD = 7.5, 67-95). **Figure 1** illustrates the MEG task, which was adapted from a prior publication (Ahveninen et al 2013). Throughout the data collection, participants were instructed to fixate on a central fixation cross.

**Figure 1:**
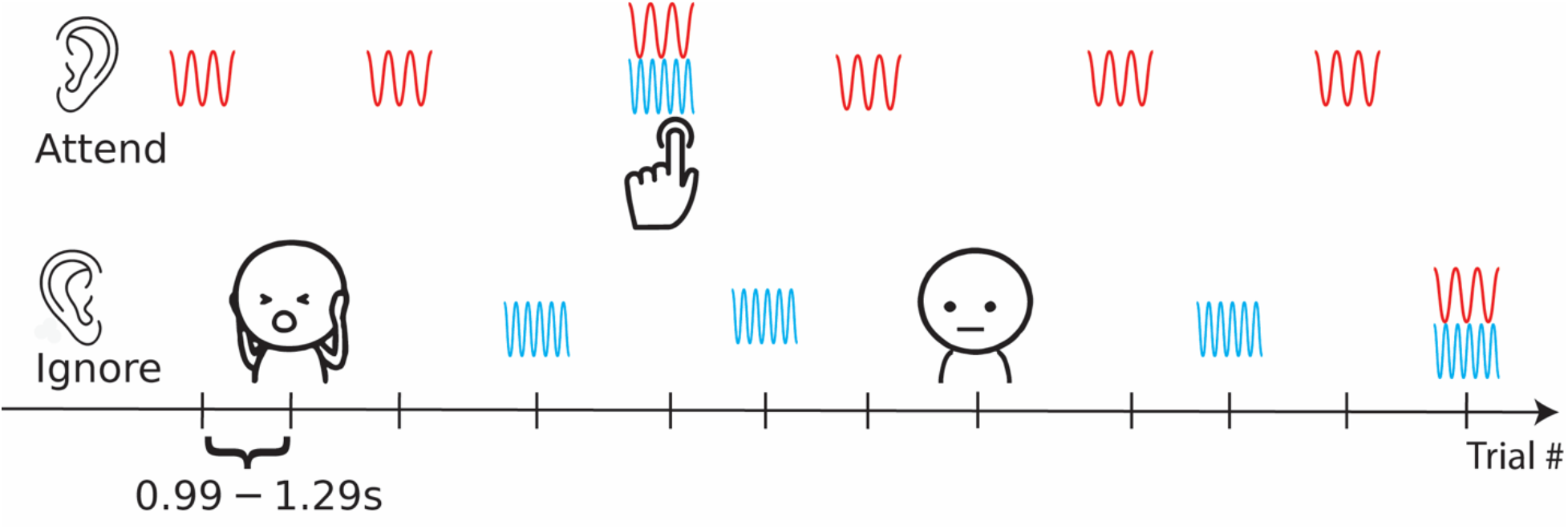
Example of the task and stimulus sequence. There were 5 different stimulus categories: (1) 800-Hz (red waveforms) or (2) 1500-Hz (blue waveforms) sinusoids lasting 0.05 s including a 0.01 s ramp-up and ramp-down, presented with intensity 65 dB and with equal likelihood across the two ears, (3) the combination of these two sounds, referred to as “target” sounds, illustrated by a red and blue waveforms, and occurring in 20% of trials in the attended (here left) ear, with intensity 68 dB, (4) complex environmental sounds shorter than 0.6 s that would be perceived as unpleasant trigger sounds, illustrated with an emoji covering ears, or (5) neutral sounds with similar parameters, illustrated here with a neutral emoji, and presented only in the unattended (“Ignore” row) ear, each with 5% probability (i.e. 5% of trials). Each run lasted for 5 minutes, where the instruction to attend versus ignore the left versus the right ear was flipped with a visual cue (not shown here) every 36 trials, at which time the stimulus streams between the left and the right ear were also flipped. There were 3 – 7 non-target sounds between target stimuli. The “target” stimuli required a button press with the left index finger for “attend left”, and right index finger for “attend right” (not analyzed as part of this study). There was a minimum of three sinusoids between trigger/non-trigger sounds. The stimulus onset asynchrony between trials was varied between 0.99 – 1.29 seconds. Illustrated here is an example where the participants were instructed to attend to the left ear (“Attend” row), with the 800-Hz sinusoid sound as the non-target sound.

The sounds in the attended ear (Fig. 1, “Attend” row) consisted of 800 Hz or 1500 Hz pure tones and target sounds that contained both the 800+1500-Hz tones. Participants were instructed to press a button as quickly as possible whenever the target sound played in the attended ear (classic dichotic oddball design). Participants were instructed to ignore all sounds in the unattended ear (“Ignore” row). Sounds in the unattended ear included the trigger and non-trigger sounds illustrated with emojis (Fig. 1, “Ignore” row), as well as 1500-Hz and 800+1500-Hz sinusoids. The present analysis focuses exclusively on the cortical responses to the trigger and non-trigger sounds that played in the unattended ear.

The trigger and non-trigger stimuli were complex environmental sounds selected from online collections of audio clips and cut to less than 0.6 s long. Before the MEG session, participants rated ten trigger sounds that typically trigger misophonia, such as chewing and coughing, on a scale of 1 (least triggering) to 10 (most triggering). Only the highest rated trigger sound, individualized for each participant, was selected to be presented during their MEG recordings. Chewing was the most common trigger sound, selected by 15 of the participants. Other sounds selected were lip-smacking (5), slurping (1), swallowing (1) and snoring (1).

Participants were also asked to rate eight potential non-trigger sounds, for example, a burst of white noise, or a clap, on a scale of 1 (least triggering) to 10 (most triggering). Any non-trigger sounds that were rated higher than 3 were removed from the set of non-trigger sounds for the participant, leaving an average of five different non-triggering sounds (range 1 – 6 non-trigger sounds) in the individualized MEG stimulus set. The order of presentation of trigger and non-trigger sounds was randomized across and within runs.

Participants were trained on the task before the MEG data were acquired, and the success rates in detecting the target tones post-training varied between ∼80% and ∼95%.

### 2.4 Structural MRI data acquisition and processing

T1-weighted magnetization-prepared rapid gradient echo (MPRAGE) structural images were acquired using a 3T Siemens Trio MRI scanner (Siemens Medical Systems, Erlangen, Germany) with a 32-channel head coil (in-plane resolution 1×1 mm; slice thickness 1.3 mm, TR 2530 ms; TI 1100 ms; TE 3.39 ms; flip angle 7°). Cortical reconstructions and parcellations were generated using the FreeSurfer software (version 7.3.3, http://surfer.nmr.mgh.harvard.edu/).

### 2.5 MEG data acquisition

The MEG data were acquired with a whole-head 306-channel neuromagnetometer (MEGIN Oy, Finland) inside a magnetically shielded room (IMEDCO, Switzerland). The 306 channels are arranged in 102 sensor triplets with two orthogonal planar gradiometers and one magnetometer. The signals were band-pass filtered at 0.1– 200 Hz prior to sampling at 1000 Hz. The position of the head was continuously recorded during the data acquisition using five head position indicator (HPI) coils attached to the scalp (Uutela et al., 2001). Locations of the HPI coils, three anatomical landmarks (nasion and pre-auricular points), and multiple additional scalp surface points were digitized using a Fastrak digitizer (Polhemus) to allow coregistering the MEG and MRI data. Additionally, electrocardiography (ECG) and electro-oculography (EOG) were recorded to detect heart activity and eye movements / blinks, respectively.

### 2.6 MEG data preprocessing

MEG data were preprocessed and analyzed using MNE-Python (Gramfort et al 2013). Noisy MEG channels were visually identified and excluded from further analyses. Temporal Signal Space Separation (Taulu & Simola 2006) was used to compensate for head movements and minimize noise from sources outside of a virtual sphere circumscribing the MEG sensors. Next, the data were bandpass filtered between 0.1 and 144 Hz, and notch filters were applied at 60 Hz and 120 Hz to remove line noise and its 1^st^ harmonic. Independent component analysis (ICA) was used to remove cardiac and oculomotor artifacts by visual inspection. The data were then epoched from -500 ms to +1500 ms with respect to sound (trigger or non-trigger) onset at time zero. Noisy epoch rejection was performed using an automated global thresholding procedure implemented in the autoreject library (Jas et al 2017). Specifically, for each condition separately, peak-to-peak amplitude rejection thresholds were estimated independently for magnetometers and gradiometers using the get_rejection_threshold function, which determines optimal thresholds via a Bayesian optimization procedure based on cross-validation across the epoched data. A 5-fold cross-validation procedure (the default) was used, with the time series down-sampled by a factor of 2 during threshold estimation to reduce computation time. Epochs in which the peak-to-peak amplitude exceeded the estimated threshold in any channel were discarded. After epoch rejection, participants contributed an average of 74.9 epochs (SD = 8.5, range 57 – 84) to the trigger condition and an average of 71.4 epochs (SD = 7.0, range 57 – 88) to the non-trigger condition. Since the number of epochs did not fulfill tests for normality, a Wilcoxon signed-rank test was used and showed that the number of accepted epochs did not significantly differ across conditions (*W* = 94.5, *p* = 0.20).

### 2.7 Source Localization

Source localization was carried out by mapping the MEG data from the planar gradiometer sensors onto cortical space using MNE-Python (Gramfort et al 2013). Source estimates were calculated on each participant’s individual anatomy, i.e., the FreeSurfer-reconstructed cortical surfaces from participant-specific T1 MRI scans. The forward solution was computed using a single-layer (inner skull) boundary element model (BEM, Hämäläinen & Sarvas 1989). Inner skull surface triangulations were generated from individual MRI structural data using the Freesurfer watershed algorithm. The cortical current distributions were obtained using the dynamic Statistical Parametric Mapping (dSPM, Dale et al 2000), with a loose orientation constraint of 0.2 and depth weighting of 0.8 (Lin et al 2006). To reduce the contribution of instrument and environmental artifacts to the MEG signals, a noise covariance matrix was obtained from the baseline of the epoched data, 200 ms before the sound onset. The noise covariance matrix was used to construct the inverse operator (Dale et al 2000, Hämäläinen & Ilmoniemi 1994). A low SNR regularization (SNR = 0.3) was applied, as is standard for induced power analyses on single-trial data using a noise covariance matrix from the pre-stimulus baseline of the epoched data.

For time-frequency domain source analyses, used for estimating power and phase-amplitude coupling, the cortical current distributions were obtained using MNE with a regularization parameter set to 0.3, a loose orientation constraint of 0.2, and a depth weighting of 0.8.

### 2.8 Regions of Interest

ROIs were defined functionally, following anatomical constraints using FreeSurfer labels. To that end, the auditory evoked response to the sinusoids (see section 2.3) were computed by epoching from -1 s to +1 s around each tone and averaging the epochs. The evoked response was then source-localized using the method described in the section above and a source time course was obtained. We then extracted the vertex in the FreeSurfer labels corresponding to the temporal transverse gyrus and the temporal transverse sulcus (*i*.*e*., Heschl’s gyrus and Heschl’s sulcus) of the Destrieux atlas (Destrieux et al 2010) which exhibited the maximum response to the auditory stimuli at 80 – 200 ms after the stimulus onsets. A circular label was then generated from this seed vertex by including all vertices within a 5 mm radius from the seed, separately in each hemisphere, thus defining the auditory ROIs for each participant. All of the analyses below were run on the source time course in this label.

### 2.9 Spectral power and phase-amplitude coupling analyses

Time-frequency representations of source-level induced power and inter-trial coherence (ITC) were computed using functions from MNE-Python (Gramfort et al 2013, Gramfort et al 2014), applying an MNE inverse solution with the above stated regularization parameter. To obtain induced power, the evoked response was first subtracted from the epochs. Frequencies of interest ranged from 4 to 120 Hz in 1-Hz steps, with the number of Morlet wavelet cycles set to f/3, yielding 1.3 cycles at 4 Hz and 40 cycles at 120 Hz, balancing a frequency-adaptive trade-off between temporal and spectral resolution. Induced power was averaged across vertices within each ROI. Baseline correction was applied using a log-ratio transform relative to a pre-stimulus baseline of −200 to 0 ms. Both induced power and ITC were averaged across vertices within each anatomical label. For this set of analyses, we focused on the frequencies 6 – 30 Hz.

Phase-amplitude coupling (PAC) was estimated at the source level using the pactools Python toolbox (Dupre la Tour et al 2017). Source time courses were extracted from the auditory cortex label using the mean operator across vertices and cropped to 0.2 – 0.6 s post-stimulus onset. The time window was chosen based on the difference between conditions in the evoked response and extended to an appropriate window length allowing for accurate estimations of the lower frequencies. Comodulograms were computed using the General Linear Model Modulation Index (GLM-MI, Penny et al 2008), with phase-providing frequencies ranging from 4 to 12 Hz (20 linearly spaced center frequencies, 1-Hz bandwidth) and amplitude-providing frequencies automatically determined from 12 Hz to 125 Hz (40 steps).

### 2.10 Statistical analyses of condition (trigger vs non-trigger) differences in evoked responses

To test for condition differences in the evoked responses, statistical comparisons between conditions were conducted using a non-parametric cluster-level permutation test as implemented in the permutation_cluster_1samp_test function in MNE-Python (Gramfort et al 2013), following the approach described by Maris and Oostenveld (Maris & Oostenveld 2007). This method controls for the multiple comparisons problem across time points by identifying contiguous temporal clusters of data points that jointly exceed a statistical threshold, rather than correcting each time point independently. Specifically, an t-statistic was computed at each time point and adjacent time points exceeding the automatically determined cluster-forming threshold — an t-threshold corresponding to p = 0.05 given the number of observations — were grouped into clusters. The sum of t-statistics within each cluster served as the cluster-level test statistic. A permutation distribution was generated by randomly reassigning observations to conditions across 1000 permutations, computing the maximum cluster-level statistic at each iteration. Cluster p-values for each observed cluster were defined as the proportion of the null distribution as extreme or more extreme than the observed cluster statistic. Clusters with p < .05 were considered statistically significant

### 2.11 Statistical analyses of condition (trigger vs non-trigger) differences in power and cross-frequency coupling

Statistical inference was performed using a non-parametric permutation test with pixel-wise correction (Maris & Oostenveld 2007). To estimate condition effects while accounting for individual differences in overall power, a mass-univariate general linear model was fit independently at each time-frequency sample, with average power as the outcome variable. The model included condition and participant as categorical predictors (power ∼ C(Participant) + C(condition), using the ordinary least squares regression implemented by the Python package statsmodels (Seabold et al, 2010), with participant included to account for between-subject variance in baseline power rather than as an effect of interest. The beta coefficient associated with the condition term at each time-frequency or frequency-frequency point was extracted and submitted to a cluster-based permutation test to correct for multiple comparisons across the time-frequency or frequency-frequency map. To build a null distribution, condition (trigger / non-trigger) labels were permuted 5000 times within each participant, preserving the within-subject data structure, and the GLM was refit on each iteration. The minimum and maximum beta values across the entire map were recorded for each permutation, forming a null distribution of extreme beta values. Significance thresholds were defined as the 2.5^th^ and 97.5^th^ percentiles of this null distribution, corresponding to a two-tailed test at α = 0.05. Observed beta values falling outside this range were considered statistically significant. Pixel-wise correction was preferred over cluster-based correction as it makes no assumptions about the spatial or temporal contiguity of effects, which was appropriate given the exploratory nature of the analyses. No individual p-values are reported, as significance is determined by whether observed beta values exceed the permutation-derived family-wise error rate threshold rather than by pointwise hypothesis testing.

### 2.12 Correlations between MEG measures and symptom severity

For each time window identified as showing a significant difference in the evoked responses to trigger versus non-trigger sounds, we extracted the peak value within the window in response to trigger and non-trigger sounds. For the clusters showing significant differences in the power and cross-frequency analyses, the mean value within the cluster was extracted for each condition. The MEG measures that showed a significant difference between the two conditions were then correlated, using Spearman’s correlation, with the relative intensity of reactions score (RIRS) from the S-Five, which gives the estimate of the intensity of reactions to triggers relative to the number of triggers.

## 3. Results

### 3.1 Evoked responses in the early auditory cortex

We began by examining the evoked responses to the trigger and non-trigger stimuli. In the left hemisphere (**Fig. 2A**), the evoked responses were characterized by a distinct peak at an average of 0.13 s post stimulus onset, followed by less pronounced deflections. There was no significant difference between the responses to the two categories of sounds. In the right hemisphere (**Fig. 2B**), the evoked responses showed two distinct peaks, the first at 0.13 s and the second at 0.25 s post stimulus onset. Cluster-based permutation analysis found significant differences between the responses to trigger and non-trigger stimuli, with the latter showing significantly larger amplitudes, first at 0.09 – 0.14 s (cluster p-value = 0.032), peaking at 0.12 s (“time window 1”), and again at 0.20 – 0.29 s (cluster p-value = 0.0034), peaking at 0.26 s (“time window 2”).

**Figure 2:**
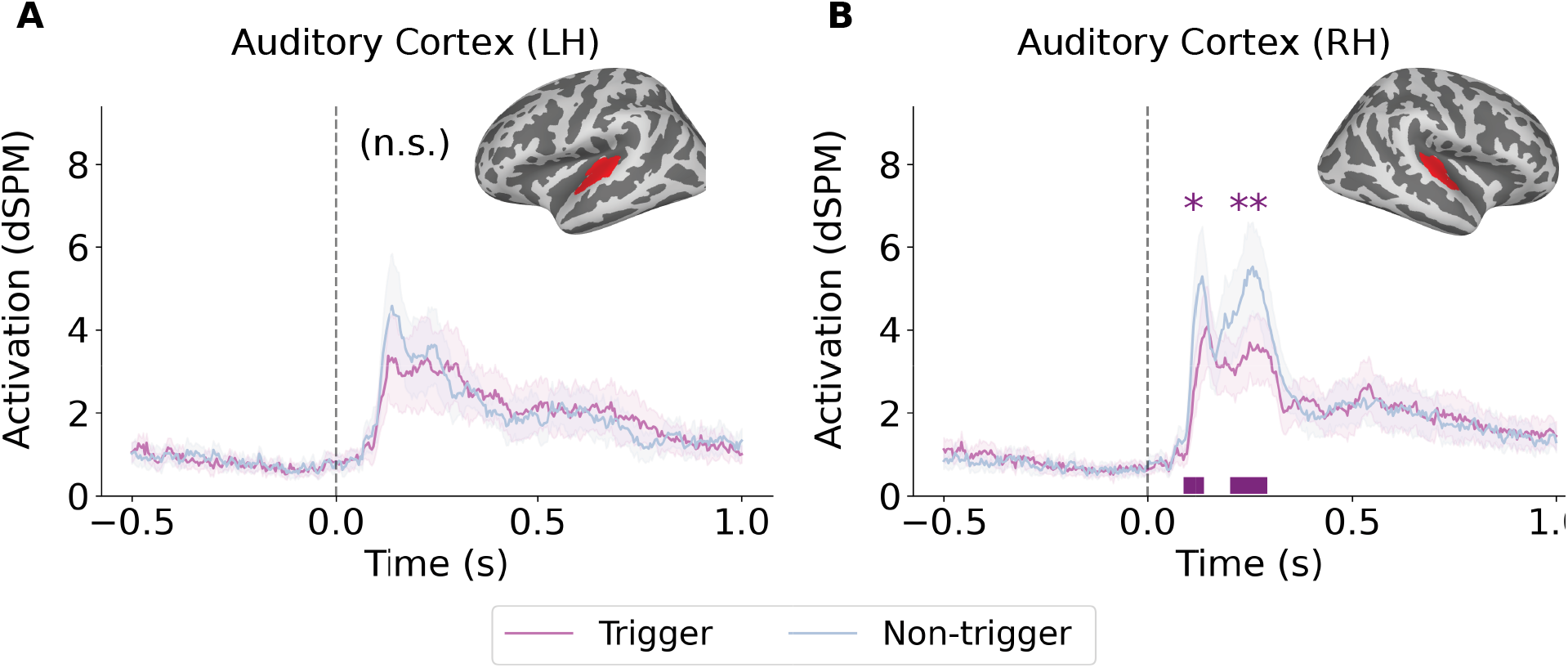
Evoked responses extracted from the auditory cortex in the left (A) and in the right hemisphere (B). In the left hemisphere there were no significant differences between the response to the trigger and non-trigger stimuli. In the right hemisphere, the responses to non-trigger stimuli were significantly stronger at two separate time windows, as marked by the horizontal purple lines at the bottom. * p < 0.05, ** p < 0.01.

### 3.2 Time-frequency oscillatory responses in the auditory cortex

Next, we examined the time-frequency (T-F) induced power maps, focusing on the time window from 0.5 s prior to stimulus onset to 1 s post stimulus onset, in the 4 **–** 30 Hz frequency band. We were primarily interested in the effect from 0 – 0.5 s after sound onset. The results for the non-trigger sounds are shown in **Fig. 3A**, and for the trigger sounds in **Fig. 3B**. For all stimuli and each hemisphere, there was an initial increase in alpha power at about 0.05 – 0.25 s, followed by a longer-lasting alpha suppression. To visualize the difference between the trigger and non-trigger sounds, we also plotted the difference T-F plot (**Fig. 3C**). Cluster-based permutation analyses showed a significant difference between conditions in each hemisphere, but only in the initial alpha increase. Significant regions in the T-F plot were defined as those exceeding the two-tailed permutation threshold (α = 0.05). The left auditory cortex showed no significant differences in the time window of interest. In the right auditory cortex, significant differences were found in the frequency range of 6 – 12 Hz between 0.11 – 0.28 s after stimulus onset. Within the cluster, to further quantify this difference, we computed the mean induced power within the significant T-F window outlined in **Fig. 3C** (right panel), for each participant in each hemisphere, and the results are shown in **Fig. 3D**. In the left hemisphere, there were no significant clusters in the time window of interest. In the right hemisphere, the mean power for trigger stimuli was 0.042 (SE = 0.015) and for non-trigger stimuli, it was 0.138 (SE = 0.015).

**Figure 3:**
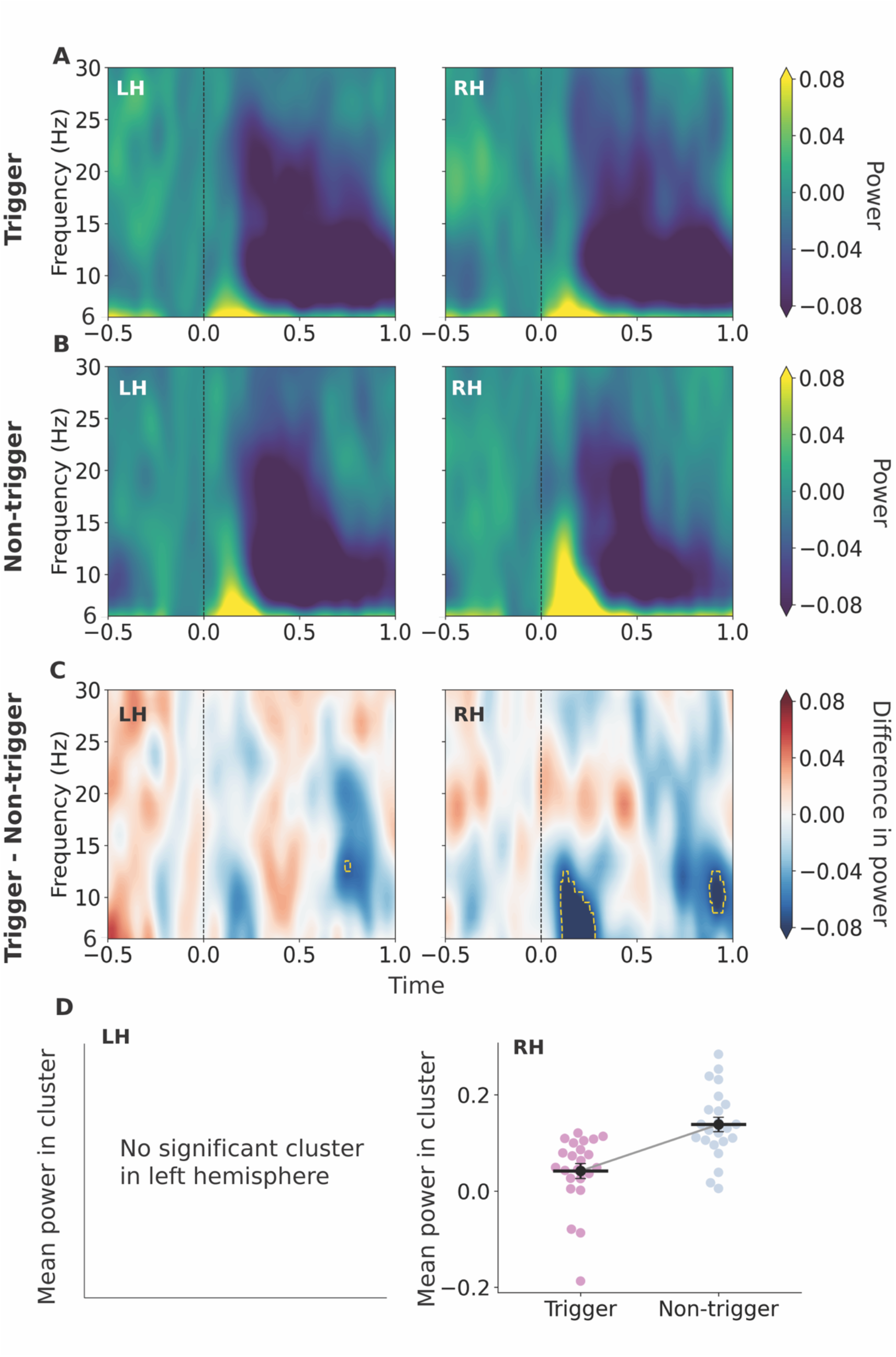
Time-frequency responses to trigger and non-trigger sounds in the auditory cortex. Raw oscillatory power (6–30 Hz) in bilateral auditory cortices shows increased theta and alpha power for both **(A)** trigger, and **(B)** nontrigger sounds. **(C)** The difference plot, trigger minus non-trigger, reveals greater theta power for non-trigger sounds in the left auditory cortex, and greater theta and alpha power in the right, with significant clusters outlined in yellow. We report the cluster within the time window of interest, which is from 0 – 0.5 s after sound onset. **(D)** Within the cluster of interest, mean power was lower for trigger than non-trigger sounds in the right hemisphere, RH: 0.042 (SE = 0.015) vs. 0.138 (SE = 0.015). There were no significant clusters in the time window of interest in the left hemisphere. LH = left hemisphere, RH = right hemisphere.

### 3.3 Stronger Alpha-Gamma Phase-Amplitude Coupling in Right Auditory Cortex for Trigger Sounds relative to non-trigger sounds

To gain a fuller understanding of the dynamics underlying the observed differences, we next computed crossfrequency coupling, and specifically phase-amplitude coupling (PAC) – the coupling between the phase of a slower frequency band and the amplitude of a faster frequency band, within the ROI, for each condition. **Fig. 4** shows the corresponding PAC results. PAC with alpha (8 – 12Hz) as the (driving) frequency for phase and beta-gamma (20 – 125 Hz) as the amplitude-frequency, is shown in **Fig. 4A** for the non-trigger stimuli, and in **Fig. 4B** for the trigger stimuli. **Fig. 4C** shows the PAC difference between two conditions, which was significant only in the right hemisphere, in two clusters, one showing alpha-beta PAC, and the other alpha-high gamma PAC, both outlined in **Fig. 4C**.

**Figure 4:**
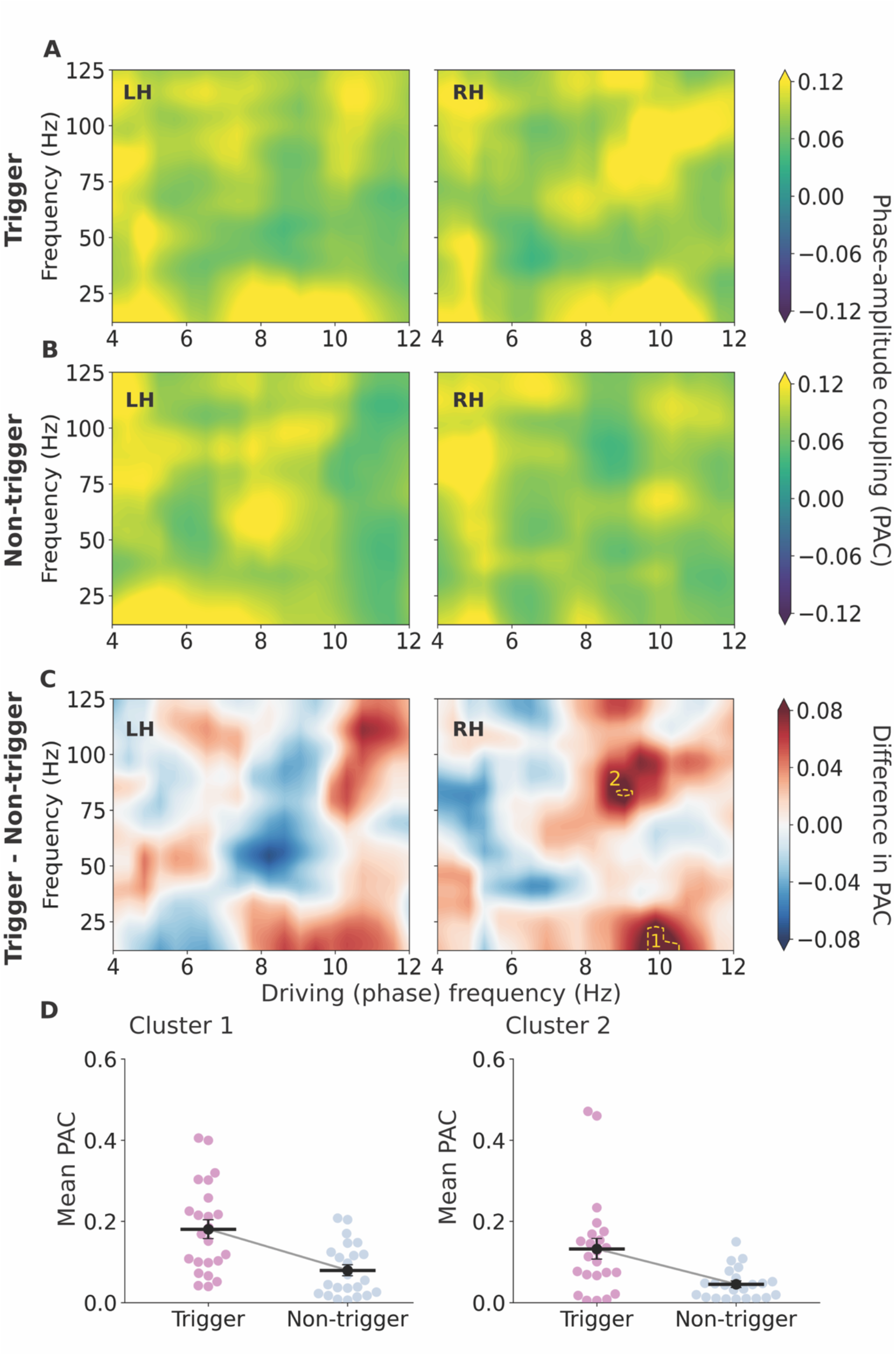
Phase-amplitude coupling (PAC) responses to trigger and non-trigger sounds in the auditory cortex. PAC between driving frequencies from 4 – 12 Hz and driven frequencies from 13 – 125 Hz in bilateral auditory cortex are shown for both **(A)** trigger and **(B)** non-trigger sounds. **(C)** The difference plot, trigger minus non-trigger, reveals greater PAC for non-trigger sounds only in the right auditory cortex, with significant clusters outlined in yellow. **(D)** Within these clusters, mean PAC was higher for trigger than non-trigger sounds in two clusters: 1, alpha-beta cluster: 0.18 (SE = 0.023) vs. 0.08 (SE = 0.013); and 2. alpha-gamma cluster: 0.13 (SE = 0.026) vs. 0.05 (SE = 0.008). LH = left hemisphere, RH = right hemisphere.

To further quantify this difference, we computed the mean PAC within these two clusters, for each participant. The results are shown in **Fig. 4D**. The mean PAC in the first cluster was 0.18 (SE=0.023) in the trigger condition and 0.08 (SE = 0.013) in the non-trigger condition. The mean PAC in the second cluster was 0.13 (SE = 0.026) in the trigger condition and 0.05 (SE = 0.008) in the non-trigger condition (Fig. 4D, right panel).

### 3.4 Correlation with misophonia symptom severity

Lastly, we tested whether any of the MEG measures that showed a significant difference between trigger and non-trigger stimuli correlated with the extent of misophonia symptoms as measured using the S-Five. To that end, we correlated the Relative Intensity of Reactions scores (RIRS) from the S-Five with each of the following MEG measures: **(1)** the peak of the amplitude of the first peak (latency = 0.09 – 0.14 s) and the second peak (latency = 0.200 – 0.292 s), of the evoked response to the trigger and non-trigger sounds in the right hemisphere (Fig. 2B), **(2)** the mean induced power in the right hemisphere in response to trigger and non-trigger sounds (Fig. 3D), and **(3)** the mean PAC in the alpha-beta cluster and the alpha-gamma cluster in the right hemisphere (Fig. 4D, right panel). **Figure 5** shows the corresponding results. Both peaks of the right hemisphere evoked response to the trigger sound correlated significantly with RIRS **(Fig. 5A** – first peak: Spearman’s rho = 0.59, *p* < 0.01, and **Fig. 5B** – second peak: Spearman’s rho = 0.43, *p* = 0.04). The mean induced power in the significant cluster found in the right hemisphere **(Fig. 5C)** did not correlate significantly with RIRS for either the trigger sound (Spearman’s rho = -0.03, *p* = 0.89) or the non-trigger sound (Spearman’s rho = 0.20, *p* = 0.37). For PAC (correlation plots not shown), there were no significant correlations with either the alpha-beta cluster (Spearman’s rho = -0.08, *p* = 0.730) or the alpha-gamma cluster (Spearman’s rho = - 0.13, *p* = 0.544).

**Figure 5:**
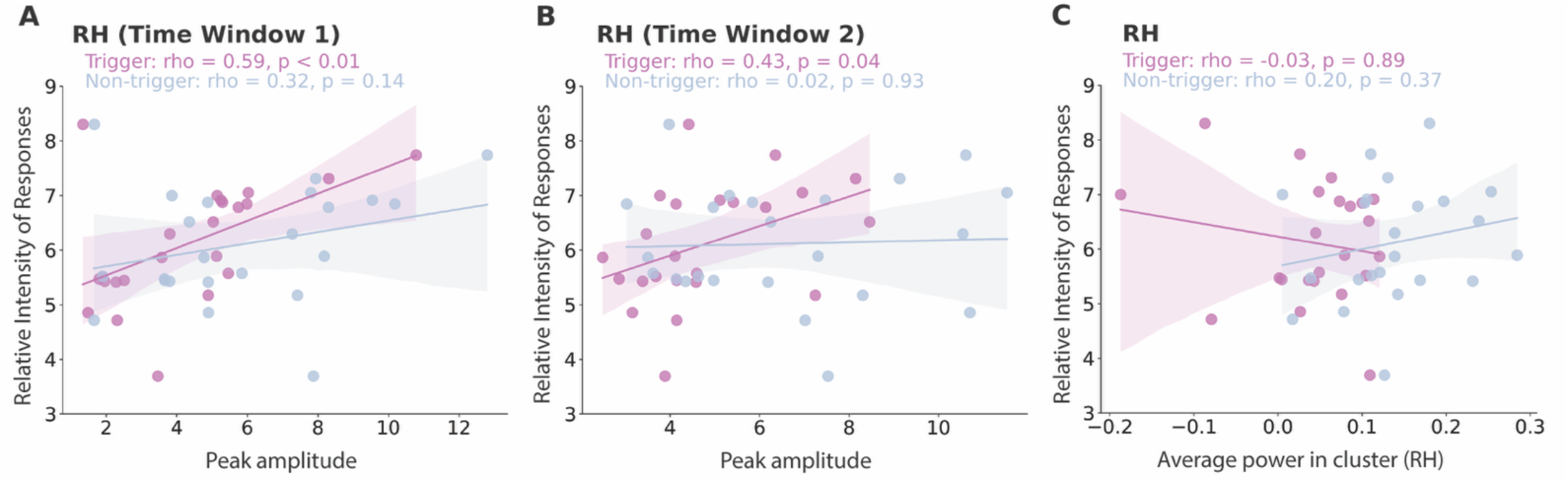
Correlations between MEG measures and symptom severity, as captured by the Relative Intensity of Response (RIRS) from the S-Five. Correlations between RIRS (higher value means more pronounced misophonia symptoms) and the evoked response peak amplitudes extracted from the right auditory cortex (*cf*. Fig. 2B) for **(A)** first peak, and **(B)** second peak. Correlations between RIRS and mean power within the cluster that showed significant difference across conditions in **(C)** the right hemisphere (*cf*. Fig. 3C RH)

## 5. Discussion

This study suggests that auditory cortex responses, measured using MEG, differ between trigger and non-trigger sounds in individuals with misophonia. Specifically, evoked responses to non-trigger sounds were more pronounced than responses to similar trigger sounds, and alpha power and phase-amplitude coupling were both different in response to trigger versus non trigger sounds. Importantly, one of the MEG measures that showed significant differences by condition also correlated with a validated behavioral measure of the magnitude of misophonia symptoms.

The identified differences in evoked responses emerged within 0.1 s of the onset of the stimulus, and first peaked on average 0.13 seconds after stimulus onset, and again at around 0.25 s. Further, the peak amplitude in response to the trigger sound in both time windows correlated positively with symptom severity, while there was no detectable correlation between symptoms severity and peak amplitude in response to the non-trigger sounds. Greater misophonia severity was not associated with a general increase in auditory cortex reactivity to all sounds, but it seems to be associated with stronger evoked responses to trigger sounds specifically. Our findings also add to the growing body of work indicating altered processing in the auditory cortex in misophonia starting as early as 0.1s or so into the cortical response. Several EEG studies have found differences in auditory evoked responses in participants with misophonia occurring at similar latencies to those noted here, specifically reduced N1 amplitudes (Karupaiah et al 2025, Schroder et al 2014) and earlier latencies of the P1, N1, and N2 peaks (Aryal & Prabhu 2024, Karupaiah et al 2025). As these prior studies have focused on responses to deviant tones, tone bursts, or consonant stimuli, our study extends these findings to ecologically valid trigger and non-trigger sounds and specifically examines these early latency responses source-localized to the auditory cortex.

The finding of less induced alpha power in auditory cortex for trigger sounds is noteworthy for several reasons. Alpha oscillations in auditory cortex have often been interpreted as reflecting inhibitory control over sensory processing, with reduced alpha power indicating release from inhibition and enhanced cortical excitability (Klimesch 2012). In typical listeners, alpha power in the auditory cortex scales with the attentional and affective significance of incoming sounds (Ahveninen et al 2017, Banerjee et al 2011, Brockhaus-Dumke et al 2008, Weisz et al 2005). It has also been shown that alpha power is increased in the auditory cortex contralateral to the ear being presented with irrelevant stimulation (Wostmann et al 2016). While in our study, both trigger and non-trigger sounds were meant to be ignored, an initial increase in alpha power was observed in response to both trigger and non-triggers sounds, but this increase in alpha was smaller for trigger sounds. In the time window of ∼0.2 s, this reduced alpha burst may reflect a failure to disengage processing of the trigger stimuli (Lombardi et al 2024). Interestingly, this alpha power difference coincides with our observed evoked response differences between trigger- and non-triggering stimuli at 0.11 – 0.28 ms, a time window in which lateral inhibition occurs (Pantev et al 2004) and GABAergic modulation of M100/M200 amplitudes takes place (Sinton et al 1986). These findings are consistent with a model where there is less inhibitory control in response to trigger sounds, which in turn aligns with prior findings that showed that trigger sounds acquire heightened salience through associative conditioning (Jastreboff & Jastreboff 2015) and extends this account to the level of early cortical dynamics.

The finding of stronger alpha-gamma cross-frequency coupling for trigger sounds reinforces this interpretation. Cross-frequency coupling, particularly PAC between alpha and gamma bands, has been proposed as a mechanism by which lower-frequency oscillations organize faster, computationally intensive gamma-band activity into coherent processing episodes (Canolty & Knight 2010). In the auditory cortex specifically, such coupling has been linked to the hierarchical encoding of complex auditory objects and to predictive coding processes that modulate the gain of auditory responses based on prior expectations (Fontolan et al 2014). In particular, the right-hemisphere finding is consistent with the well-established right dominance for the processing of complex non-speech sounds and for prosodic and affective auditory processing (Zatorre 2001, Zatorre & Belin 2001). In misophonia, the stronger alpha-gamma PAC might reflect a state of heightened and temporally organized cortical excitability when trigger sounds are encountered, whereby the auditory cortex is not merely disinhibited (as reflected by alpha suppression) but enters a dynamically reorganized oscillatory state that may amplify and sustain the cortical representation of trigger stimuli. This interpretation is also consistent with computational models suggesting that PAC facilitates the selective amplification of task-relevant sensory representations (Hyafil et al 2015).

More generally, in combination, our findings align with and extend the emerging neurobiological literature on misophonia showing divergent cortical responses to trigger versus non-trigger stimuli. It has been shown, using fMRI, that activation in the superior temporal cortex in response to audiovisual trigger stimuli differed from the response to non-trigger stimuli (Schröder et al 2019). Additionally, several fMRI studies consistently identified heightened amygdala activation and aberrant connectivity between auditory cortex and limbic structures as hallmarks of misophonia trigger processing (Eijsker et al 2019, Kumar et al 2017). Kumar and colleagues additionally reported increased myelination of connections between auditory cortex and emotion-processing areas in misophonia, suggesting a structural correlate for aberrant auditory-affective coupling. More recent work using resting-state fMRI has implicated the anterior insula and salience network in misophonia, consistent with the prominent interoceptive and visceral quality of trigger responses (Eijsker et al 2019). Our findings complement these observations, by extending the differentiation of responses into early stages of the cortical auditory processing hierarchy.

Some limitations of the current study warrant acknowledgment. First, our sample was relatively small, and replication in larger cohorts is needed to establish the robustness of these oscillatory signatures, and test whether significant differences might in fact be present more extensively in left hemisphere as well. Second, while the study benefited from the advantage of each participant being their own control by comparing across conditions rather than between groups, the study did not include a matched non-misophonia control group, and future work would ideally add a comparison with matched controls. Furthermore, the study focused only on early responses, originating in or near areas lower in the auditory cortices hierarchy. Future studies are needed that extend these and similar analyses to other cortical areas. Lastly, it is worth noting that an exploration of the results with the full cohort of 40 participants, including the ones with the erroneous sound volume normalization, showed similar results but with lower significance values, most likely as a result of the unnormalized sound volumes.

In conclusion, the present study demonstrates that processing of trigger versus non-trigger sounds is significantly altered in misophonia, especially in the right auditory cortex, as evidenced by differences in the amplitude of the evoked response, lower level of induced alpha, and stronger alpha-gamma PAC in response to trigger sounds. Importantly, the amplitude of the evoked response to the trigger sounds correlated with behaviorally assessed extent of misophonia symptoms. Mapping the neural correlates of misophonia is important not only for improving our understanding of the neural substrates underlying the condition, but also for the development and assessment of targeted interventions (Brout et al 2018, Neacsiu et al 2022). These findings advance our understanding of the neural basis of misophonia by revealing a dynamic neural signature of trigger processing in the auditory cortex following stimulus onset. More broadly, these results support a view of misophonia in which aversive sound-specific reactions emerge, at least in part, from altered neural dynamics at early stages of sensory processing, with implications for the development of neuromodulatory and psychophysiological interventions targeting auditory cortex excitability.

